# Genomic Signatures of Domestication in a Fungus Obligately Farmed by Leafcutter Ants

**DOI:** 10.1101/2024.07.02.601708

**Authors:** Caio A. Leal-Dutra, Joel Vizueta, Tobias Baril, Pepijn W. Kooij, Asta Rødsgaard-Jørgensen, Benjamin H. Schantz-Conlon, Daniel Croll, Jonathan Z. Shik

**Affiliations:** Section for Ecology and Evolution, Department of Biology, University of Copenhagen, Universitetsparken 15, 2100 Copenhagen, Denmark; Laboratory of Evolutionary Genetics, Institute of Biology, University of Neuchatel, Rue Emile-Argand 11, CH-2000, Neuchâtel, Switzerland; Department of General and Applied Biology, São Paulo State University (UNESP), Institute of Biosciences, Rio Claro, SP, Brazil; Smithsonian Tropical Research Institute, Apartado 0843-03092, Balboa, Ancon, Republic of Panama

## Abstract

The naturally selected fungal crop (*Leucoagaricus gongylophorus*) farmed by leafcutter ants shows striking parallels with artificially selected plant crops domesticated by humans (e.g., polyploidy, engorged nutritional rewards, dependence on cultivation). To date, poorly resolved *L. gongylophorus* genomes based on short-read sequencing have constrained hypotheses about how millions of years under cultivation by ants shaped the fungal crop genome and potentially drove domestication. We use PacBio HiFi sequencing of *L. gongylophorus* from the leafcutter ant *Atta colombica* to identify 18 putatively novel biosynthetic gene clusters that likely cemented life as a cultivar (e.g., plant fragment degradation, ant-farmer communication, antimicrobial defense). Comparative analyses with cultivated and free-living fungi showed genomic signatures of stepwise domestication transitions: 1) free-living to ant-cultivated: loss of genes conferring stress response and detoxification, 2) hyphal food to engorged nutritional rewards: expansions of genes governing cellular homeostasis, carbohydrate metabolism, and siderophore biosynthesis, and 3) detrital provisioning to freshly cut plant fragments: gene expansions promoting cell wall biosynthesis, fatty acid metabolism, and DNA repair. Comparisons across *L. gongylophorus* fungi farmed by three leafcutter ant species highlight genomic signatures of exclusively vertical clonal propagation and widespread transposable element activity. These results show how natural selection can shape domesticated cultivar genomes towards long-term ecological resilience of farming systems that have thrived across millennia.

## Introduction

Crop domestication involves coevolutionary processes where selectively propagated individuals are actively planted and cultivated in protected growth environments (Purugganan, 2022). Across generations, the stringent direction selection imposed by farmers can yield crop adaptations that promote productivity and nutritional optimization while also causing genomic changes that preclude a free-living existence (Araki et al., 2007). Domesticated genomes can thus differ markedly from those of free-living ancestors, with fixation of advantageous alleles (Tenaillon et al., 2023), reduced genetic diversity (Fornasiero et al., 2022), epigenetic changes (Cao et al., 2023) and polyploidy (Wang et al., 2022). While domesticated genome evolution is usually studied in the plant crops that humans artificially selected over the past 10,000 years, there also exist lineages of fungi obligately farmed—and potentially domesticated—by insects over millions of years (Mueller et al. 2005). We propose that these independent fungicultural origins can provide important insights into whether and how natural selection can drive crop domestication.

Consider the fungus-farming ‘attine’ ants (tribe: Attini) that emerged as the planet’s earliest known farmers over 60 million years ago in the rainforests of South America (Branstetter et al., 2017). In small colonies of perhaps tens to hundreds of workers, the attine ants adopted a lineage of Basidiomycete fungus that they provisioned with tiny bits of detritus (e.g., insect frass), protected from environmental stress, consumed for food, and vertically transmitted to daughter colonies (Schultz & Brady, 2008). These early farming lineages are represented today by the palaeoattine and lower neoattine ants whose fungi generally provide their farmers with undifferentiated hyphae as the food source (Möller, 1893; Mueller et al., 2001; Mueller et al., 2005). As the attines spread across the Americas, their fungal cultivars acquired domesticated traits that coincided with stepwise transitions towards incrementally greater farm size and complexity in the higher neoattine clade with up to thousands of workers and the crown group leafcutter ants with up to millions of workers (Shik et al., 2014; Branstetter et al., 2017).

All leafcutter ants are assumed to farm the fungal species *Leucoagaricus gongylophorus* and we hypothesized that its unique genomic adaptations set the stage for the genus *Atta* to become ecologically important herbivores across diverse ecosystems from southern Argentina to northern Texas (Mueller et al., 2011; Branstetter et al., 2017). These adaptations are expressed most prominently in gongylidia (Aylward et al., 2013; De Fine Licht et al., 2014), the swollen hyphal cells that are otherwise unique in the fungal kingdom and provide both nutritional rewards for ant farmers (Martin et al., 1969; Quinlan & Cherrett, 1979; Bolander et al., 2023; Leal-Dutra et al., 2023) and enzymes that ants vector unharmed in fecal droplets to promote degradation and detoxification of provisioned plant material (De Fine Licht et al., 2010; De Fine Licht et al., 2013; Schiøtt & Boomsma, 2021). To date, explorations of genomic changes underlying such domesticated traits have been constrained because we have lacked high-resolution genomic resources to delimit syntenic regions and, genetic and structural variations potentially responsible for enabling *L. gongylophorus* to convert plant material that ants cannot eat to a nutritionally optimized and readily digestible diet.

This knowledge gap owes mainly to the genetic complexities of the *L. gongylophorus* because each cell contains up to 17 heterokaryotic nuclei (Kooij et al., 2015; Carlson et al., 2017). This functional polyploidy combined with limited accuracy of existing genomes have precluded studies of changes in genome synteny as well as structural variation within haplotypes. For instance, the short-read *L. gongylophorus* genome of Aylward et al. (2013) isolated from a Panamanian colony of *Atta cephalotes* provides the most current and best cited for genome for this fungal species (Floudas et al., 2015; Dentinger et al., 2016; Nygaard et al., 2016; Guo et al., 2023; Martiarena et al., 2023). While the Aylward et al. (2013) genome was the state of the art at the time of its publication, it provides an incomplete and fragmented assembly that cannot delimit key features of genomic change. At an even more basic level, various analytical methods do not even agree on the size of the *L. gongylophorus* genome (Kooij & Pellicer, 2020). Case in point, a second short-read-based assembly of *L. gongylophorus* from a Mexican *Atta mexicana* colony (Accession: GCA_022457215) was recently deposited on GenBank, but it yielded an assembly 1.5 times larger (152 Mb) than the Aylward et al. (2013) assembly (101 Mb).

Moreover, transposable elements (TEs) are thought to promote domestication in human domesticated crops since they can generate useful genomic novelty through altered coding sequences, chromosome rearrangements, and modified gene regulatory networks (Bourque et al., 2018; Thieme & Bucher, 2018; Muszewska et al., 2019). It was thus intriguing that Clutterbuck (2017) used the Aylward et al. (2013) assembly to suggest that the *L. gongylophorus* genome is also enriched in transposable elements (TEs) due to high levels of CpG depletion. Yet, this TE hypothesis remains inconclusive because the sequence data generated by Aylward et al. (2013) do not span complete repeat regions, and thus incorrectly resolve the TE landscape by collapsing highly repetitive regions. Current genomic inference thus relies on sequencing data that are too fragmented and incomplete to accurately delimit genomic complexities arisen during the domestication process.

In the present study, we aim to resolve the issues of assembly fragmentation and completeness using PacBio HiFi long-read sequencing to produce a high-quality, annotated genome assembly of *L. gongylophorus* from a Panamanian colony of the leafcutter ant, *Atta colombica*. We then harnessed this assembled genome to investigate four hypotheses: 1) the genome of *L. gongylophorus* is enriched with TEs; 2) the ancestral shift towards a mutualistic lifestyle farmed by ant colonies was facilitated by the adaptive evolution or exaptive elaboration of novel biosynthetic gene clusters (BGC); 3) the major domestication transitions were marked by high levels of gene gain and loss; 4) the lack of gene flow and sexual recombination within and across populations of vertically transmitted *L. gongylophorus* clones generates divergent genomic lineages that are highly polymorphic and structurally variable.

## Results

### High-quality genome assembly

Our long-read genome assembly of *L. gongylophorus* (Ac2012-1) yielded a more contiguous and complete assembly than earlier genomes (Table 1) and enabled a more robust estimate of genome size. From ca. 206,000 HiFi reads (2.47 Gbp) of up to 50 kb length, we generated an initial draft genome of 187.09 Mb with 1217 contigs (maximum length of 7.91 Mb), 34.51% of GC content, and an N50 of 2.18 Mb (Suppl. File 1). To reduce the duplication level likely caused by the heterozygosity and polyploidy, we collapsed redundant contigs using Redundans (see methods) to create a final consensus assembly that we used in the downstream analyses. This final assembly was 74.31 Mb with 39 scaffolds (40 contigs; maximum length of 7.91 Mb), 37.32% of GC content, and a N50 of 6.12 Mb. The genome completeness analysis of the final Ac2012-1 assembly using BUSCO found 89.6% complete Agaricales orthologs, of which 88.1% were single copy and 1.5% were duplicated (Table 1). This assembly exceeded the completeness of the previously assembled cultivars of *A. cephalotes* (73.8% complete Agaricales orthologs) and *A. mexicana* (61.6% complete Agaricales orthologs).

**Table 1:**
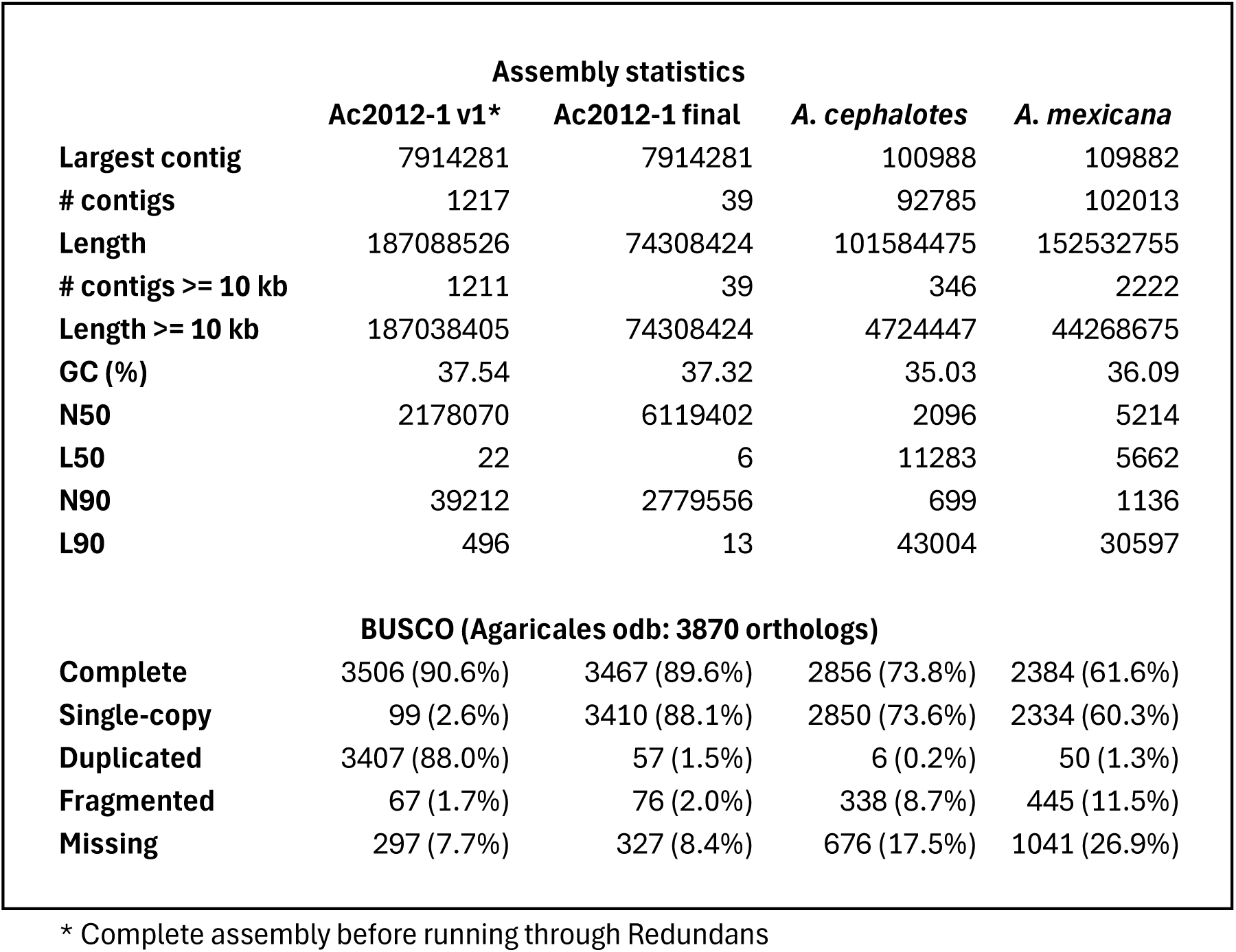
Detailed assembly statistics and BUSCO results for the three *L. gongylophorus* assemblies used in this study.

### Extensive transposable element abundance in the L. gongylophorus genome

Our results support the hypothesis of a high TE content as our long-read genome of *L. gongylophorus* has a total TE content of 67.78% of total genome size (50.36Mb), with 633 distinct TE classifications. The genome is dominated by retrotransposons, with LTR elements and LINEs comprising 32.3% (24.01Mb) and 13.1% (9.73Mb) of total genome size respectively (Suppl. Fig. S1, Suppl. Table S1, Suppl. File 2). Unclassified elements comprise 22.2% of total genome size (16.53Mb), which can be partially attributed to the general lack of TE surveys in the fungi kingdom. There are few DNA repeats (0.1%), with a complete absence of helitrons (also termed rolling circles) and SINEs, although one family of each was found in the pre-Redundans assembly (Suppl. Fig. S1, Suppl. File 2).

### Functional annotation highlights novel metabolic insights

Analysis with the Funannotate pipeline (Palmer & Stajich, 2022) showed that our long-read assembly contained 8,298 gene models, representing 85 tRNA and 8,213 coding sequences (CDS) from which 7,511 were assigned with at least one functional trait. These gene models provided the basis for further analyses that identified syntenic genes forming biosynthetic gene clusters using the fungiSMASH webserver with AntiSMASH v.7.1.0 (Blin et al., 2023).

From this, we detected 20 BGCs, including two terpenes with high similarity to (+)-δ-cadinol, and 18 that are putatively novel (53 in the pre-redundans assembly; see Suppl. File 3). Within these BGCs we identified five terpenes, five NRPS-like, three fungal-RiPP-like, two indoles, two T1PKS, and one NI-siderophore. The secondary metabolites produced by these BGCs likely have diverse functions, including communication, antimicrobial activity, and iron uptake (Bingle et al., 1999; Carroll & Moore, 2018; Kües et al., 2018). In contrast, the same analysis performed on the two short-read *L. gongylophorus* assemblies provided far lower resolution, revealing only five gene clusters (*A. cephalotes* isolate) or one cluster (*A. mexicana* isolate) (Suppl. File 3). Therefore, our assembly improves the resolution of BGC inference and highlights novel molecular mechanisms and evolutionary processes possibly involved in fungal domestication by leafcutter ants.

### Linking domestication transitions and genome evolution

We first reconstructed a consensus phylogenomic tree that resolved relationships between our assembly and those of seven closely related fungal genomes which were publicly available on NCBI database (Fig. 1, Suppl. Table S2). We based the tree on the alignment of 527,465 amino acids from 1136 BUSCO orthologs (of 3870 total) that were present as complete and single-copy in all genomes analysed. As expected, this analysis recovered the three *L. gongylophorus* isolates as a sister clade to *Leucoagaricus* sp. from a colony of the higher neoattine *Trachymyrmex arizonensis* (Fig. 1). The tree also indicates that attine-farmed fungi are polyphyletic, supporting previous phylogenies inferred from only a single protein encoding gene plus four ribosomal markers (Mehdiabadi et al., 2012; Mueller et al., 2018). Specifically, a clade with two free-living *Leucoagaricus* species (*L. leucothites* and *L. birnbaumii*) separates the fungal mutualist of the lower neoattine *Cyphomyrmex costatus* from the other fungal mutualists (Fig. 1).

**Figure 1:**
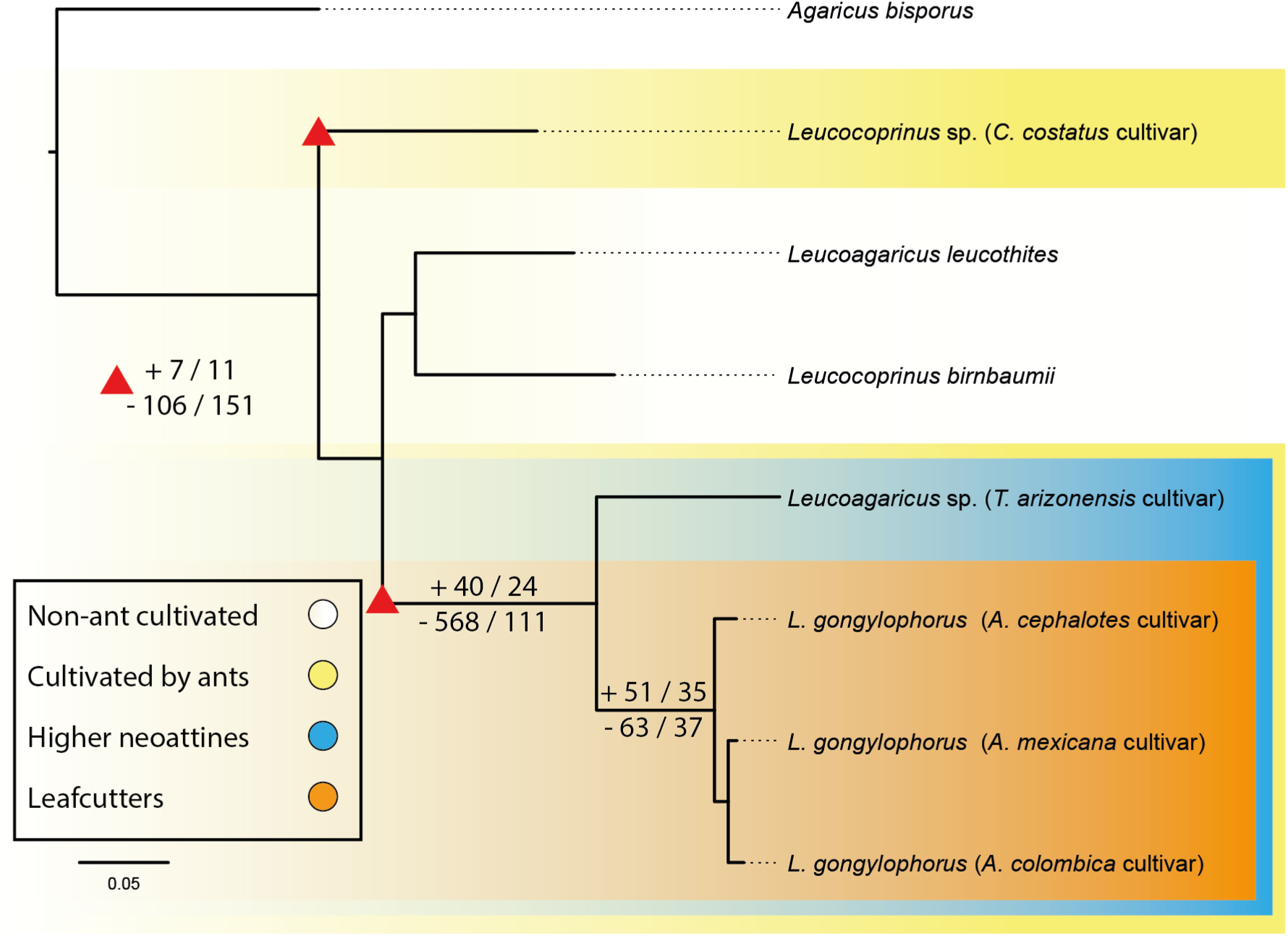
Maximum-likelihood phylogeny showing domestication transitions in ant-farmed and closely related free-living fungi. The tree was reconstructed from 1136 sequences of amino-acid from single-copy orthologs shared by all samples. Domestication transitions indicated by colored boxes: 1) yellow: from free-living to ant-cultivated (polyphyletic), 2) blue: from hyphal food to engorged nutritional rewards (higher neoattines transition), and 3) orange: from detrital provisioning to freshly cut plant fragments (leafcutters transition). Gene gain / expansion (**+**), and gene loss / contraction (**-**) are represented for each of these transitions (indicated by the red triangle for the transition to ant-cultivated species). All branches received maximum UFBoot support (100%). Scale bar: amino acid substitutions per site.

We next performed a gene gain and loss analysis to explore genetic modifications during three domestication transitions: 1) from free-living to ant-cultivated, 2) from hyphal food to engorged nutritional rewards in the higher neoattine clade, and 3) from detrital provisioning to plant fragment provisioning in the leafcutter clade (Fig. 1).

The transition to ant cultivation coincided with substantial gene losses relative to non-ant cultivated fungal clades since 106 orthogroups were lost and 151 gene families were reduced, while only 7 orthogroups were gained and only 11 gene families were expanded (Fig. 1). The domestication transition to specialized nutritional rewards (higher neoattine-Leafcutter clade vs. all other clades) coincided with further genetic losses, with 568 orthogroups lost and 111 gene families being reduced, and with only 40 orthogroups being gained and 24 gene families expanded. In contrast, the transition to fresh-vegetation provision by leafcutters was associated with less-pronounced genomic changes since similar numbers of orthogroups were lost (63) and gained (51), and similar numbers of gene families were reduced (37) and expanded (35).

We next explored which functions were gained (or expanded) and lost (or contracted) across each domestication transition by performing a functional enrichment analysis within these orthologs (Fig. 2; Suppl. File 4). Some genomic trends were consistent across all ant-farmed fungi, including the loss or contraction of genes conferring stress response and detoxification and the gain of genes for macroautophagy and transposition of TEs (Fig 2A). Subsequent genetic gains or expansion were specific to either the transition to higher neoattines (cellular homeostasis, carbohydrate metabolism, siderophore biosynthetic pathway; Fig. 2B) and the transition to leafcutters (cell wall biosynthesis and organization, fatty acid metabolism, and several processes for DNA repair; Fig. 2C). Losses or contraction of gene functions showed similar stepwise changes as the higher neoattine fungal clade lost genes related to carbohydrate-mediated transcription and signaling (Fig. 2B), and the leafcutter fungal clade lost genes related to demethylation, catabolizing certain proteins and macromolecules, and countering osmotic stress (Fig. 2C). These incremental and directional functional gene increases and reductions thus appear to reflect the novel selection pressures encountered across successive domestication transitions.

**Figure 2:**
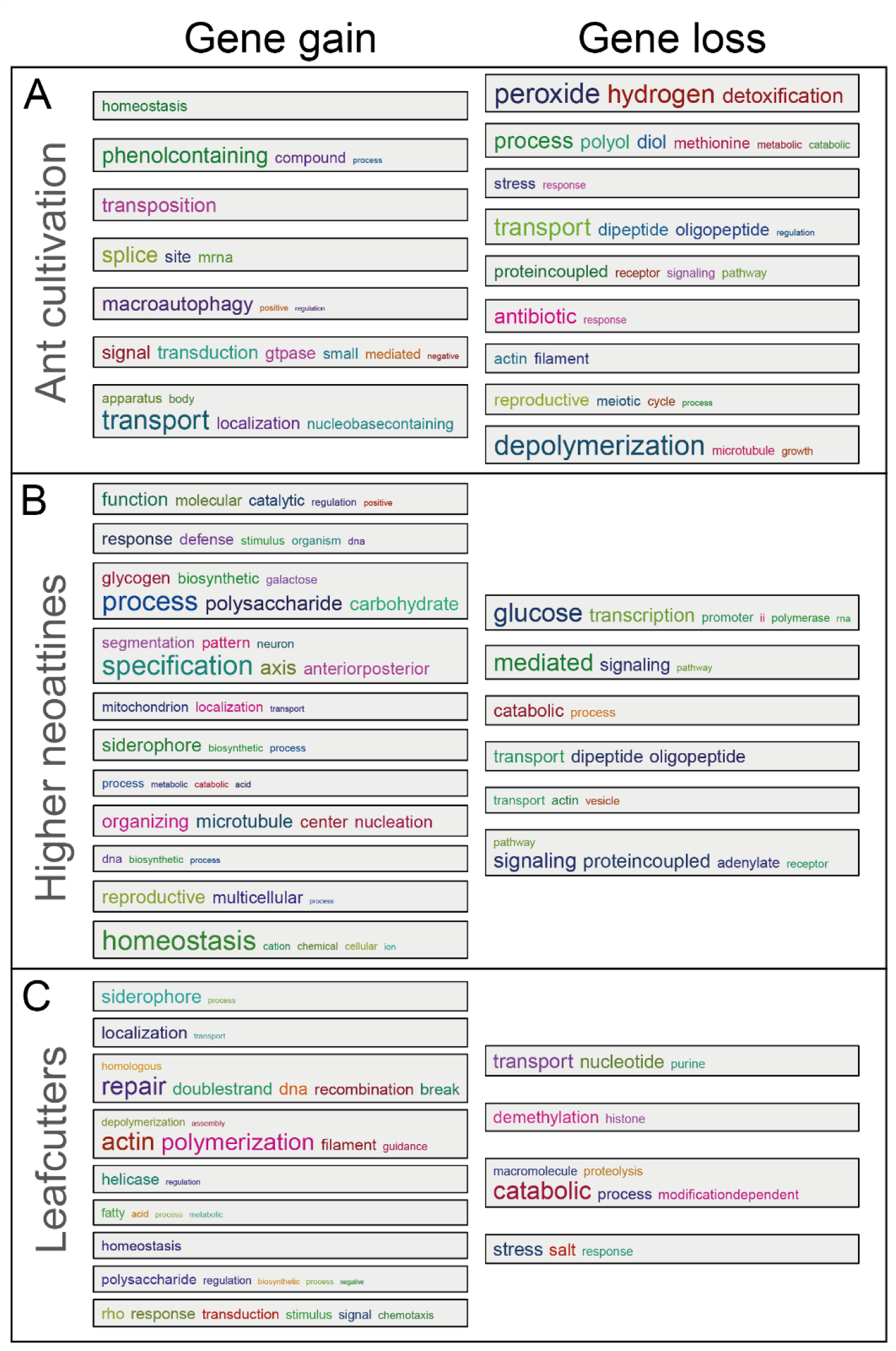
Word clouds representing the clusters of enriched Biological Process terms from the gene ontology enrichment analysis. Each box clusters significant GO terms from similarity matrices of functional terms corresponding to gene gain (or gene family expansion; left column) and loss (or gene family contraction; right column) through each step of the domestication process noted in Figure 1.

### Genome evolution under vertically transmitted propagation

While all leafcutter ants are assumed to farm a single fungal species, we tested whether this lumping assumption may belie genomic divergence among reproductively isolated strains of *L. gongylophorus* cultivated by different leafcutter species. To quantify pairwise differences between the three available *L. gongylophorus* assemblies, we first used a k-mer analysis by calculating the Jaccard containment index (JCI) of k-mer composition for each assembly. This analysis indicates that our *L. gongylophorus* assembly shares only ca. 35.9% of its k-mers with the assembly of the same fungus cultivar species isolated from of *A. mexicana*, and even fewer k-mers (ca. 8.4%) with the assembly when isolated from *A. cephalotes* (Fig. 3A). The genomes thus exhibit extensive divergence, reflecting the genetic distance of these vertically transmitted, and thus reproductively isolated, but conspecific fungal lineages.

**Figure 3:**
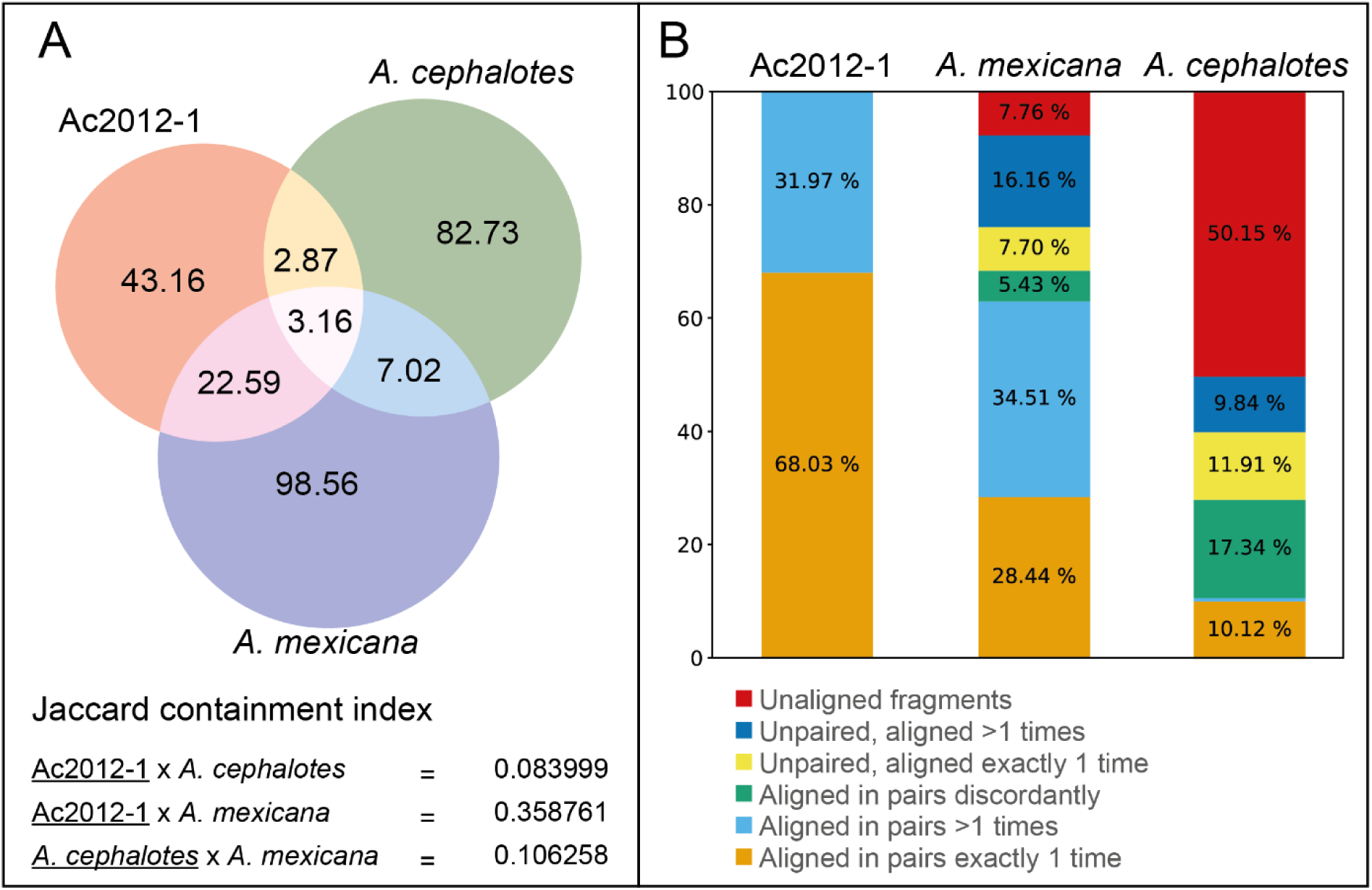
Genomic similarity comparisons of *L. gongylophorus* assemblies across leafcutter ant species. **(A) Sourmash k-mer Analysis:** The Venn diagram displays the number of k-mer (k=51) MinHashes shared between three *L. gongylophorus* assemblies, in millions of k-mer MinHashes, alongside their Jaccard Containment Indexes (JCI), with the shorter assembly underlined. This analysis indicates that approximately 8.4% and 35.9% of the Ac2012-1 k-mers are found in the genomes of *A. cephalotes* and *A. mexicana*, respectively. The *A. mexicana* cultivar contains approximately 10.6% of k-mers from *A. cephalotes*. **(B) Short-Read Simulation (SRS) Analysis:** The bar chart presents the mapping rate of simulated read fragments from Ac2012-1 aligned to itself and the other reference assemblies. The percentage of fragments aligning to each assembly is color-coded to indicate paired and unpaired alignments, as well as non-alignments. These results suggest large structural variations and a high level of divergence among the assemblies. They also highlight a closer genomic relationship between Ac2012-1 and *A. mexicana* cultivar, reflected by a higher alignment rate, in contrast to the more divergent *A. cephalotes* cultivar.

We note that k-mer analyses may overstate levels of evolutionary distance because they conservatively discard all mismatches arising from single or short nucleotide variations such as single nucleotide polymorphisms (SNPs) or insertions or deletions (indels). We thus developed a less conservative ‘short-read simulation’ (SRS) approach that fragments each assembly to simulate paired-end 150 bp read sequences (PE150) that can then be tested for levels of correspondence when pairwise mapped to each other assembly (*See Methods for more detailed explanation*). The SRS test yielded 496,972 fragments from our assembly, 492,852 from the *A. cephalotes* cultivar assembly, and 800,852 from the *A. mexicana* cultivar assembly (Table 2).

**Table 2:** Mapping rate of fragments generated in the SRS analysis.

| **Simulated PE150 fragments from Ac2012-1 from *A. colombica* (total fragments 496,972)** |  |  |  |
| --- | --- | --- | --- |
| Assemblies | *A. colombica* | *A. cephalotes* | *A. mexicana* |
| Overall fragments alignment rate | 100.00% | 49.85% | 92.24% |
| Fragments aligned in pairs exactly 1 time | 68.03% | 10.12% | 28.44% |
| Fragments aligned in pairs >1 times (duplicated) | 31.97% | 0.68% | 34.51% |
| Fragments aligned in pairs discordantly | 0.00% | 17.34% | 5.43% |
| Fragments (unpaired) aligned exactly 1 time | 0.00% | 11.91% | 7.70% |
| Fragments (unpaired) aligned >1 times (duplicated) | 0.00% | 9.84% | 16.16% |
| Unaligned fragments | 0.00% | 50.15% | 7.76% |
| **Simulated PE150 fragments from *A. cephalotes* cultivar (total fragments 492,852)** |  |  |  |
| Assemblies | *A. colombica* | *A. cephalotes* | *A. mexicana* |
| Overall fragments alignment rate | 34.87% | 100% | 38.61% |
| Fragments aligned in pairs exactly 1 time | 10.45% | 84.62% | 8.60% |
| Fragments aligned in pairs >1 times (duplicated) | 0.93% | 15.38% | 5.59% |
| Fragments aligned in pairs discordantly | 15.54% | 0.00% | 9.20% |
| Fragments (unpaired) aligned exactly 1 time | 4.13% | 0.00% | 5.83% |
| Fragments (unpaired) aligned >1 times (duplicated) | 3.84% | 0.00% | 9.39% |
| Unaligned fragments | 64.97% | 0.00% | 61.39% |
| **Simulated PE150 fragments *A. mexicana* cultivar (total fragments 800,852)** |  |  |  |
| Assemblies | *A. colombica* | *A. cephalotes* | *A. mexicana* |
| Overall fragments alignment rate | 76.28% | 46.18% | 100% |
| Fragments aligned in pairs exactly 1 time | 31.00% | 7.91% | 27.60% |
| Fragments aligned in pairs >1 times (duplicated) | 9.79% | 0.67% | 72.40% |
| Fragments aligned in pairs discordantly | 13.58% | 12.81% | 0.00% |
| Fragments (unpaired) aligned exactly 1 time | 8.35% | 13.40% | 0.00% |
| Fragments (unpaired) aligned >1 times (duplicated) | 13.56% | 11.38% | 0.00% |
| Unaligned fragments | 23.72% | 53.82% | 0.00% |

We first mapped the fragments of our assembly Ac2012-1 against itself to establish a baseline correspondence level (Fig. 3B). While all the fragments mapped successfully, approximately 32% fragments mapped both ‘in pairs’ and more than once, possibly reflecting that the genome contains large numbers of TEs and duplicated regions. When we next mapped Ac2012-1 fragments to each other assembly, ca. 7.8% failed to map to the *A. mexicana* assembly and 50.15% failed to map to the *A. cephalotes* assembly. This suggests a high divergence between the assemblies. Additionally, 29.3% of the Ac2012-1 fragment pairs aligned discordantly (i.e., both fragments of the pair aligned but not as a pair) or aligned ‘unpaired’ (i.e., only one fragment of the pair) to the *A. mexicana* cultivar, and 39.1% showed the same result when mapped to for the *A. cephalotes* cultivar. This suggests high levels of structural variation (e.g., transposition, duplications, inversions). Thus, despite high overall levels of genomic divergence, our assembly from *A. colombica* shares more similarities with the geographically distant cultivar of *A. mexicana* than with the cultivar of *A. cephalote*s from the same study locality in Panama.

## Discussion

The complex multinucleate genome of *L. gongylophorus* has previously constrained explorations of whether and how continuous directional selection by leafcutter ant farmers has produced a domesticated cultivar. Such inference is of special interest in this fungus because it exhibits a suite of traits like polyploidy, elaborated nutritional rewards, and obligate dependence on cultivation also seen in plant crops domesticated by humans. In this study, we used PacBio HiFi long-read sequencing to overcome previous analytical issues and produce a highly contiguous megabase-length assembly that opens new perspectives in synteny, transposable elements, and clusters of co-regulated genes and genetic rearrangements across successive domestication transitions. We are further able to confirm the hypothesis that genomic rearrangements such as inversions, duplications, and transpositions set the stage for genetic novelty potentially driving fungal domestication. Moreover, since 90% of our consensus assembly is comprised of only 13 contigs, this suggests chromosome-length resolution comparable to the other highest quality available fungal assemblies (e.g. button mushroom *Agaricus bisporus*) (Sonnenberg et al., 2016).

### Transposable elements provide opportunities for domesticated trait evolution

Transposable elements represented around two thirds of the *L. gongylophorus* genome, which is extremely high for a fungus (Clutterbuck, 2017; Lebreton et al., 2022). Specifically, 67% of the total TE content was composed of LTRs and LINEs, which have shown high levels of activity throughout the evolutionary history of *L. gongylophorus* but a notable deficiency in recent activity (Suppl. Fig. S1). This supports an earlier study by Clutterbuck (2017) that indirectly inferred abundant TEs based on CpG depletion levels that are known to silence and immobilize TEs (Gladyshev, 2017). Which is further supported by the loss of genes related to demethylation, increasing the chances of CpG transitions and fixation of old TEs. This is important because high activity of TEs can set the stage for rapid genome evolution (Raffaele & Kamoun, 2012) which has been seen in other domesticated crops of humans (Cai et al., 2022). Abundant TEs are also known in fungal ectomycorrhiza symbioses (Hess et al., 2014; Lebreton et al., 2022).

Given the expansion of genes related to transposition in the first domestication transition (Fig. 2), we further predict that abundant TEs may be a universal feature of all attine cultivated fungi. Thus, TEs may serve a dual role of promoting genomic variation and driving coevolutionary processes in fungal symbioses. This perspective strengthens the role of TEs in shaping the genomes of symbiotic partners, enhancing the resilience of the symbiosis and allowing them to thrive in fluctuating environments.

### Functional consequences of genome evolution

The annotation made possible by our contiguous assembly not only enhances our understanding of the *L. gongylophorus* genome, but it also enables exploration of potentially novel BGCs and their yet-to-be discovered roles in mutualisms. Specifically, we identified and annotated 20 BGCs (18 putatively new) with diverse functions. For example, terpenes and indoles are known to mediate fungal communication from the cellular level to interspecific signaling (Kües et al., 2018), potentially enabling the fungus not only to repel fungal competitors or pathogens but also to precisely align its physiological needs with those of the ants. Additionally, some terpenes like (+)-δ-cadinol, and several NRPS, and T1PKS have antifungal and antimicrobial properties (Bingle et al., 1999; Felnagle et al., 2008; Ringel et al., 2022; Vinha et al., 2023). Consequently, these BGCs produce secondary metabolites that could potentially act in colony defense. Furthermore, NI-siderophores may facilitate iron acquisition, essential for Fenton reaction in *L. gongylophorus* fungus gardens known from another leafcutter genus *Acromyrmex* (Schiøtt & Boomsma, 2021), allowing the initial breakdown of biomass even when iron is made unavailable by plant-secreted iron chelators (Dellagi et al., 2005; Carroll & Moore, 2018). Our high-quality genome assemblies can thus help test hypotheses about evolution of domestication by investigating how metabolic changes potentially favor adaptation and specialization of domesticated symbionts.

### Stepwise genomic evolution across domestication transitions

A specific gene set likely required for the ancestral cultivars be recruited for cultivation by ants shared by all extent cultivars and absent in the free-living relatives. In this first step of the domestication process we highlight the loss of detoxification genes in the cultivars, which could be attributed to the ants’ preliminary screening for toxic compounds in the substrate (Herz et al., 2008), which reduces the selective pressure on the fungi to retain these genes. These results also support the hypothesis that autophagic repurposing of cellular components is important not only for gongylidium formation (Leal-Dutra et al., 2023), but likely to other mechanisms in the symbiosis, since autophagy-related genes were gained or expanded across all cultivars.

The evolution of gongylidia as hyphal swellings containing metabolites with multimodal functions was a decisive domestication event in the fungal lineage farmed by higher neoattine and leafcutter ants (Mueller et al., 2001; Schultz & Brady, 2008). Gongylidia have clear analogies with enlarged fruits of domesticated plant crops farmed by humans, both for their morphology and nutrient content that would likely be maladaptive in a free-living state, and for their polyploid genomic architecture (Alam & Purugganan, 2024). Since gongylidia likely ended an avenue for a free-living existence, it is perhaps not surprising that this domestication transition incurred the greatest amount of gene loss (568) we observed. Additionally, among the few gained and expanded gene functions, cellular homeostasis and carbohydrate metabolism might have favored the nutritional stability of gongylidia, while siderophore biosynthesis could support acquisition of iron for Fenton reaction as mentioned before. The loss of genes related to carbohydrate-mediated transcription and signaling may also have also facilitated gongylidium engorgement since they reduce the cellular response to sugar accumulation (Koch, 1996; Jiang et al., 1997) and allow sugar to concentrate without a cellular feedback response.

The *L. gongylophorus* fungus enables the leafcutters to access an herbivorous niche that is otherwise unique among the ants (Hölldobler & Wilson, 2011; Shik et al., 2021) and our results help delimit the genomic adaptations that could have enabled this domestication transition. For instance, *L. gongylophorus* is enriched with genes related to cell wall biosynthesis and modification that are upregulated during gongylidium formation (De Fine Licht et al., 2014). Fatty acid metabolism genes are also enriched in ways that may produce metabolites that shape ant farming behaviors as exemplified by Khadempour et al. (2021). *L. gongylophorus* is also enriched for DNA repair genes that may be a response to genomic instability caused by the loss of oxidative stress regulatory genes that occurred at the onset of ant cultivation.

### Genomic consequences of obligate vertical crop transmission

Despite similarities in *L. gongylophorus* assemblies observed in the phylogeny using protein encoding genes (Fig. 1), we observed considerable variation at the whole genome level. This variation likely emanates from the reproductive isolation of these fungi since they are propagated vertically as clonal inoculates carried by dispersing foundress queens (Weber, 1972; Hölldobler & Wilson, 2011; Kooij et al., 2015). This propagation mode has likely promoted founder effects among distinct fungal populations that accumulate lineage-specific variation (e.g. SNPs, structural variants, indels). While such variation is expected when comparing allopatric samples from Panama and Mexico, we also observe variation in *L. gongylophorus* farmed sympatrically by two different species of *Atta* leafcutter ants. These results strongly support a hypothesis that colonies of different leafcutter ant species are strong barriers to genetic admixture. Further research and the generation of additional high-quality genomes of the attine-cultivated fungi will be crucial to understand the genome evolution patterns across these ant-domesticated fungus.

In conclusion, we use long-read sequencing of the *L. gongylophorus* genome to reveal with novel resolution the intricate genetic changes that accompany each step of fungal domestication, from the initial suitability for ant cultivation to advanced modifications that enabled large-scale leafcutter farming systems. Our findings underscore that fungal genome modifications (e.g., gene gain-loss, TE-driven rearrangements) were crucial for expanding the ecological niche and long-term evolutionary stability of leafcutter ants. Moreover, our results provide a robust genomic framework for further research on how domestication can evolve.

## Methods

### Sample acquisition

We generated genome sequencing data using *L. gongylophorus* fungus sampled from an *Atta colombica* colony (Ac2012-1) collected from Soberanía National Park, Panama and maintained at the University of Copenhagen under controlled conditions (25°C, 70% RH, minimal daylight). We isolated the fungus in axenic cultures in 90-mm diameter Petri dishes with 20-ml potato dextrose agar (PDA) and kept it in the dark at 25°C. To extract DNA, we optimized fungal biomass production by introducing the PDA-cultivated fungus into 250 ml of potato dextrose broth (PDB). This was agitated at 140 rpm in the dark at 25°C. After 30 days, the mycelia were sieved using a nylon mesh, rinsed with sterile distilled water, and freeze-dried for 48 hours.

### Available genomic data

To evaluate the sequence and structure of our genome assembly, we used two publicly available short-read genome assemblies of *L. gongylophorus* from *Atta cephalotes* (Aylward et al., 2013) and *Atta mexicana* (unpublished, but see Vigueras et al. (2017) for details) colonies for comparisons. We further included five other genome assemblies to perform a gene gain and loss analysis across the phylogeny: one fungal cultivar from a lower neoattine, *C. costatus* (Nygaard et al., 2016), one fungal cultivar from a non-leafcutter higher neoattine, *T. arizonensis* (Beigel et al., 2021), and three from non-ant cultivated fungal species, *A. bisporus* (Morin et al., 2012), *L. birnbaumii* (unpublished, GenBank GCA_027627405), and *L. leucothites* (Floudas et al., 2020). Details about each of these additional genome assemblies are provided in Suppl. Table S1.

### DNA Extraction and Sequencing

High molecular weight DNA was extracted from fungal tissue using the CTAB-based lysis and precipitation protocol of Jones and Schwessinger (2021). Briefly, about 150 mg of freeze-dried mycelia were ground in liquid nitrogen and resuspended in 10 mL of lysis buffer (4% CTAB, 1% PVP-40, 100 mM Tris, 20 mM EDTA, 1.2 M NaCl, pH 8) along with RNAse A and proteinase K, both at 200 µg per mL. The mixture was placed in a rotating mixer at 10 rpm and incubated at 60 °C for 90 min. These samples were then purified using phenol:chloroform:isoamyl alcohol (25:24:1) followed by an additional purification step with an equal volume of chloroform:isoamyl alcohol (24:1). The DNA was then precipitated by incubating the supernatant with two volumes of precipitation buffer (2% CTAB, 100 mM Tris, 20 mM EDTA, pH 8) and rotating at 60 °C for 2.5 h. This mix was centrifuged, and the resulting DNA pellet washed with 80% ethanol and centrifuged again. After air drying, the pellet was resuspended in 150 µL of 10 mM Tris-HCL (pH 8). All centrifugation steps were performed at the speed and temperature guidelines provided by Jones and Schwessinger (2021).

Extracted samples were sent for long-read PacBio HiFi sequencing at the Beijing Genomics Institute (BGI, Hong-Kong, China). A PacBio SMRTbell library was prepared using the SMRTbell Template Prep Kit 2.0 and Sequel II Binding 2.2 (PacBio, CA, USA), and loaded onto the sequencer by diffusion. The sequencing was conducted on a Sequel II platform (PacBio, CA, USA), employing a Sequel II Sequencing Kit 2.0 and a SMRT Cell 8M Tray, and managed by the instrument control software and the Primary SW (both v.10.1.0.119549). HiFi reads were generated using the *Circular Consensus Sequencing* workflow (CCS) v.6.4.0 (https://github.com/PacificBiosciences/ccs) with default parameters. Sequencing statistics were performed using Samtools v.1.16 (Danecek et al., 2021).

### Genome Assembly

An initial genome assembly was performed using the HiCanu algorithm of Canu software v.2.2 (Koren et al., 2017; Nurk et al., 2020) with default settings and genome size of 40 Mbp, as previously reported by (Kooij & Pellicer, 2020). Assembly quality was assessed using QUAST v.5.2.0 (Gurevich et al., 2013), BUSCO v.5.4.5 (Manni et al., 2021) with the Agaricales OrthoDB v10 (Kriventseva et al., 2018) as reference.

We next used Redundans v.1.01 (Pryszcz & Gabaldón, 2016) to reduce the redundancy of our assembly, that generates polymorphic duplicated contigs due to heterozygosity. After testing multiple parameter combinations and assessing the resulting assemblies using BUSCO— while assigning arbitrary weights to single-copy versus duplicates, completeness, and assembly size (Suppl. Table S3)—we set Redundans to collapse shorter contigs with over 82% similarity and a minimum of 75% overlap with longer contigs. With the final assembly, we identified and removed mitochondrial contigs using MitoFinder v.1.4.1 (Allio et al., 2020).

### Transposable elements prediction and annotation

Next, we curated and annotated transposable elements in *L. gongylophorus* using the Earl Grey TE annotation pipeline (v.3.0) with default parameters (Baril et al., 2022; Baril et al., 2023). Briefly, de novo consensus sequences were initially identified and subsequently refined in Earl Grey using a “BLAST, Extract, Extend, Trim” process (Platt et al., 2016). Following final annotation, Earl Grey processes TE loci to remove overlapping annotations and to defragment annotations likely originating from the same TE insertion event. The TE consensus library and GFF annotation file are provided in Suppl. File 2.

### Functional annotation

The Ac2012-1 genome annotation was performed using the Funannotate pipeline v.1.8.13 aided by the transcriptome and RNAseq reads from the same fungus isolate published in previous study (Leal-Dutra & Shik, 2023; Leal-Dutra et al., 2023). This process was done with default Funannotate settings It incorporated several ab-initio gene prediction training tools, including Augustus v3.3.1 (Stanke et al., 2008), CodingQuarry v1.3(Testa et al., 2015), GeneMark-ES/ET v4.71 (Lomsadze et al., 2014), glimmerHMM v3.0.4 (Majoros et al., 2004), Snap v2006-07-28 (Korf, 2004) and tRNAscan-SE v2.0.11 (Chan et al., 2021). These tools utilized splicing information generated by PASA v2.3.3 (Haas et al., 2003). Finally, EVidenceModeler v1.1.1 (Haas et al., 2008) was used to generate consensus gene models. For further functional annotations, Funannotate employed InterProScan v5.55-88.0 (Jones et al., 2014), EggNOG Mapper v2.1.1 (Cantalapiedra et al., 2021), AntiSMASH v7.0.1 (Blin et al., 2023), SignalP v5.0b (Almagro Armenteros et al., 2019), Phobius v1.1 (Käll et al., 2004), and DIAMOND v2.1.6.160 (Buchfink et al., 2021). Functional predictions were assigned by matches to the CAZy (Lombard et al., 2013), InterPro (Paysan-Lafosse et al., 2022), MEROPS (Rawlings et al., 2014), Pfam (Finn et al., 2013), and Swiss-Prot databases (The UniProt Consortium, 2022).

The final assembly and annotation of *Leucoagaricus gongylophorus* Ac2012-1, were deposited at NCBI GenBank under the BioProject accession PRJNA879936. Assembly and annotation of the pre-Redundans version of this genome can be found as Supplementary File 1.

### Phylogeny reconstruction

We produced a phylogenomic tree to confirm the placement of our Ac2012-1 assembly within the public genomes using an amino acid dataset from 1136 single-copy orthologs shared by the assemblies. We identified these orthologs using BUSCO as described above. To prepare the dataset, we first created individual fasta files for each ortholog and aligned the sequences using MUSCLE v.3.8.425 (Edgar, 2004) with default parameters. Next we trimmed each alignment with trimAl v.1.4.rev15 (Capella-Gutiérrez et al., 2009) using a heuristic selection to decide the optimal method for trimming (setting option -automated1). All the trimmed alignments were concatenated, and a RAxML-style partition file was created. Finally, we used IQ-TREE v.2.1.3 (Minh et al., 2020) to reconstruct a maximum-likelihood phylogenomic tree, first using the built-in ModelFinder (Kalyaanamoorthy et al., 2017) allowing the program to merge partitions to increase model fit followed by tree inference with 1000 ultrafast bootstraps (UFBoot) (Hoang et al., 2018).

### Gene gain/loss and gene family expansion/contraction analyses

To analyze gene gain or loss and gene family expansion or contraction, we first used the phylogeny reconstructed in this study to delimit the clades into three distinct groups, each representing a step in the domestication transition (Fig. 1). We then performed the analyses for each of these groups. First, we looked at the transition from free-living to cultivation by ants, comparing non-ant cultivated fungi with all cultivars (including the cultivar of the lower neoattine *C. costatus* and the clade of the higher neoattines that includes the cultivar of *T. arizonensis* and the three *L. gongylophorus*; yellow box in Fig. 1). Then we focused on the transition from hyphal food to engorged nutritional rewards, comparing the clade of the higher neoattines cultivars (blue box in Fig. 1) with the free-living and lower attine cultivar. And finally, we tested the transition from detrital provisioning to freshly cut plant fragments, comparing only the *L. gongylophorus* clade (orange box in Fig. 1) with the remaining samples.

We used the protein sequences predicted in all the studied assemblies to assess the number of clade-specific genes across the phylogeny. First, we used Orthofinder v2.5.4 (Emms & Kelly, 2019) to infer orthology and set the program to search for sequences using ultra-sensitive DIAMOND, split paralogous clades, and infer maximum-likelihood trees from multiple sequence alignments. The resulting phylogenetic hierarchical orthogroups were further explored to categorize them into single-copy orthogroups (those with 1 copy in at least 75% of the genomes), multi-copy orthogroups (one or more copies in at least 75% of the genomes, and at least 25% of the genome with 2 or more copies), and taxon-specific orthogroups (present in a single clade or assembly). We also functionally annotated each orthogroup using InterProScan and eggNOG-mapper, setting the filter threshold of bit score to 40, and percentage of identity, query cover and subject cover to 20%.

We used the inferred orthogroups to retrieve those that are associated with each group of clades, representing different steps in the domestication process. We specifically performed a strict gain/loss analysis to identify group-specific orthogroups that comprise: 1) putatively new genes that are only present among all genomes in each group and absent in the other clades, and 2) putatively lost genes that are absent in each group, but present in all the other clades. We also identified gene family expansion and contraction by comparing the number of gene copies present in the species in each group against the remaining clades. We finally performed a gene ontology enrichment analysis using GO gene sets with biological process (BP) ontology within various orthogroups to highlight the overrepresented functions in genes gained or expanded, and lost or reduced across the three domestication steps using R package TopGO (Alexa & Rahnenfuhrer, 2023). To visualize these enrichments, we used simplifyEnrichment (Gu & Hübschmann, 2023) to cluster the significant GO terms with similarity matrices of functional terms, and produce a word cloud with the common BP ontology shared by genes within these clusters. It is important to note that although we used a yeast database to assign GO terms to the orthogroups, several terms are derived from animal studies, reflecting the current gaps in our understanding of Agaricales gene functions. Hence, these genes should be further investigated in experimental studies.

### Assessing sequence similarity across L. gongylophorus assemblies

To evaluate the genomic similarities among the three *L. gongylophorus* assemblies, we employed a two-pronged approach. First, a stringent Jaccard Containment-based k-mer composition analysis, and second, a more relaxed alignment-based analysis. This dual methodology was chosen to provide a comprehensive and nuanced understanding of the genomic similarities.

The first approach, using the Jaccard Containment-Based k-mer composition analysis, was conducted with Sourmash v.4.8.2 (Brown & Irber, 2016). In this method, k-mers (k=51) were generated from each genome assembly to create MinHash sketches. These sketches were then utilized to calculate similarity using the Jaccard containment index. This index is particularly effective for comparing genomes of varying quality and size, as it considers the proportion of shared k-mers in relation to the size of the smaller genome set (Pierce et al., 2019). This provides a robust similarity measure, accommodating the differences in the genome assemblies’ quality. In this analysis, pairwise containment scores, which ranged from 0 to 1, were compiled for further evaluation. A score of 1 signifies a high degree of similarity, indicating that all 51-mers in the smaller genome were found in the larger genome.

However, considering the unique characteristics of *L. gongylophorus*, which is both polyploid and highly heterozygotic, the k-mer-based approach has limitations. It restricts similarity searches to exact k-mer matches, lacking the capacity to account for nucleotide-level variations such as base substitutions. To address this, we developed a more lenient method that assesses sequence similarities while ignoring small sequence variations. This involved aligning small fragments from each assembly to the other *L. gongylophorus* assemblies. To circumvent potential misalignments of large contigs that might occur using whole-genome alignment tools, and due to the absence of short reads associated with these assemblies, we evaluated assembly similarities by simulating paired-end 150 bp reads (PE150) derived from each genome. For this, each contig was segmented into 600 bp fragments, further divided into two 450 bp inserts with a 300 bp overlapping region. The initial and final 150 bp of each insert were allocated to the left and right reads files, respectively, with the final 150 bp reverse complemented (Suppl. Fig. S2). We then used Bowtie2 v.2.5.0 (Langmead & Salzberg, 2012) to align these short sequences with the assemblies using the default parameters. Since Bowtie2 utilizes an alignment scoring system, which accommodates nucleotide-level variations (e.g. SNPs) and structural variations (e.g. indels, duplications, inversions, etc.), this method provided a more relaxed comparison, capturing lineage-specific mutations.

Therefore, by using both the stringent k-mer based approach and a more relaxed alignment-based analysis we benefit from their complementary strengths. The k-mer based approach provides a precise and stringent measure of similarity, especially useful when dealing with genomes of different sizes and quality. On the other hand, the alignment-based analysis offers a broader, more inclusive assessment, capturing more general similarities that the stringent method might overlook. Together, these methods enable a more holistic and accurate assessment of the genomic similarities among the *L. gongylophorus* assemblies.

## Acknowledgements

Authors are grateful for the useful comments and discussions provided by Jacobus Koos Boomsma, Søren Rosendahl and Gareth W. Griffith. General laboratory assistance was provided Sylvia Mathiasen and Rasmus Stenbak Larsen. Data processing and analysis were performed using Computerome, the National Life Science Supercomputer at DTU (www.computerome.dk). The Ministerio de Ambiente, Republica de Panama provided permits for colony collection and sample exportation (SEX/A-31-12). STRI provided fieldwork assistance and permits requests support.

