## Supplementary table 1 for "Genomic Signatures of Domestication in a Fungus Obligately Farmed by Leafcutter Ants"

**Table S1:** Transposable elements quantification for pre-Redundans and final assembly.

| | Ac2012-1 pre-Redundans | | | Ac2012-1 final version | | |
|---|---|---|---|---|---|---|
| | Number of TEs | Lenght sum | Genome % | Number of TEs | Lenght sum | Genome % |
| LTR | 235 | 116906 | 0.06 | 0 | 0 | 0.00 |
| LTR/Caulimovirus | 1385 | 570004 | 0.30 | 0 | 0 | 0.00 |
| LTR/Copia | 9473 | 4450949 | 2.38 | 2616 | 1413088 | 1.90 |
| LTR/Gypsy | 86664 | 65372488 | 34.94 | 21924 | 22596053 | 30.41 |
| LINE | 565 | 521771 | 0.28 | 178 | 170607 | 0.23 |
| LINE/I-Jockey | 129 | 81701 | 0.04 | 128 | 101338 | 0.14 |
| LINE/Penelope | 205 | 165382 | 0.09 | 93 | 74900 | 0.10 |
| LINE/Tad1 | 17866 | 20872089 | 11.16 | 6875 | 9379407 | 12.62 |
| LINE/L2 | 148 | 141023 | 0.08 | 0 | 0 | 0.00 |
| LINE/R1 | 87 | 96618 | 0.05 | 0 | 0 | 0.00 |
| DNA/Kolobok-H | 63 | 94555 | 0.05 | 47 | 71033 | 0.10 |
| DNA/Dada | 139 | 60037 | 0.03 | 0 | 0 | 0.00 |
| RC/Helitron | 35 | 11133 | 0.01 | 0 | 0 | 0.00 |
| SINE/5S-Sauria-RTE | 76 | 162335 | 0.09 | 0 | 0 | 0.00 |
| Unknown | 86554 | 32056936 | 17.13 | 46295 | 16531954 | 22.25 |
| Low_complexity | 68 | 10089 | 0.01 | 29 | 4640 | 0.01 |
| Simple_repeat | 247 | 42746 | 0.02 | 121 | 22284 | 0.03 |
| **Total** | **203939** | **124826762** | **66.72** | **78306** | **50365304** | **67.78** |
