## Supplementary table 2 for "Genomic Signatures of Domestication in a Fungus Obligately Farmed by Leafcutter Ants"

**Table S2:** Detailed information about available genomic data used in this study.

| Assembly | Host-ant species | Sample | Sample locality | Sequencing technology | Accession | Reference |
|---|---|---|---|---|---|---|
| *A. colombica* | *Atta colombica* | Ac2012 | Gamboa, Panama | PacBio | PRJNA879936 | This study |
| *A. cephalotes* | *Atta cephalotes* | Ac12 | Gamboa, Panama | 454 pyrosequencing | GCA_000382605.1 | Aylward et al., 2013 |
| *A. mexicana* | *Atta mexicana* | LEU18496 | Coatepec, Veracruz, Mexico | Illumina + 454 | GCA_022457215.2 | Vigueras et al., 2017 |
| *T. arizonensis* | *Trachymirmex arizonensis* | KB180720.3 | Southwestern Research Station, Arizona, USA | PacBio | GCA_019327925.1 | Beigel et al., 2021 |
| *C. costatus* | *Cyphomyrmex costatus* | SymC.cos | Gamboa, Panama | Illumina | GCA_001563735.1 | Nygaard et al., 2016) |
| *Leucocoprinus birnbaumii* | Non-ant cultivated | VT141 | Cuc Phuong National Park, Ninh Binh, Vietnam | IonTorrent | GCA_027627405.1 | Unpublished |
| *Leucoagaricus leucothites* | Non-ant cultivated | CBS 146.42 | Sweden | PacBio | GCA_013368445.1 | Floudas et al., 2020 |
| *Agaricus bisporus* | Non-ant cultivated | H97 | unknown | Illumina + 454 | GCA_000300575.2 | Morin et al., 2012 |
