## Supplementary table 3 for "Genomic Signatures of Domestication in a Fungus Obligately Farmed by Leafcutter Ants"

**Table S3:** BUSCO results from  redundans test runs. Results in each column are represented in a colour gradients from green (best) to red (worst).

| Redundans params | Complete | Single | Duplicate | Fragmented | missing | # Scaffolds | # Contigs | Length |
|---|---|---|---|---|---|---|---|---|
| id = 80, ovlp = 70 (a) | 88.90% | 88.40% | 0.50% | 2.00% | 9.10% | 35 | 36 | 71708553 |
| id = 81, ovlp = 70 (a) | 88.90% | 88.40% | 0.50% | 2.00% | 9.10% | 35 | 36 | 71708553 |
| id = 82, ovlp = 65 (b) | 89.60% | 88.10% | 1.50% | 2.00% | 8.40% | 40 | 41 | 74558126 |
| id = 82, ovlp = 70 (b) | 89.60% | 88.10% | 1.50% | 2.00% | 8.40% | 40 | 41 | 74558126 |
| id = 82, ovlp = 75 (b) | 89.60% | 88.10% | 1.50% | 2.00% | 8.40% | 40 | 41 | 74558126 |
| id = 82, ovlp = 80 | 89.60% | 88.10% | 1.50% | 2.00% | 8.40% | 41 | 42 | 74614353 |
| id = 83, ovlp = 60 (c) | 89.60% | 87.70% | 1.90% | 1.90% | 8.50% | 43 | 44 | 75022751 |
| id = 83, ovlp = 65 (c) | 89.60% | 87.70% | 1.90% | 1.90% | 8.50% | 43 | 44 | 75022751 |
| id = 83, ovlp = 70 (c) | 89.60% | 87.70% | 1.90% | 1.90% | 8.50% | 43 | 44 | 75022751 |
| id = 83, ovlp = 75 (d) | 89.60% | 87.70% | 1.90% | 1.90% | 8.50% | 44 | 45 | 75078978 |
| id = 83, ovlp = 80 (d) | 89.60% | 87.70% | 1.90% | 1.90% | 8.50% | 44 | 45 | 75078978 |
| id = 85, ovlp = 60 (e) | 89.60% | 84.50% | 5.10% | 1.90% | 8.50% | 53 | 54 | 77701617 |
| id = 85, ovlp = 65 (e) | 89.60% | 84.50% | 5.10% | 1.90% | 8.50% | 53 | 54 | 77701617 |
| id = 85, ovlp = 70 (e) | 89.60% | 84.50% | 5.10% | 1.90% | 8.50% | 53 | 54 | 77701617 |
| id = 85, ovlp = 80 (e) | 89.60% | 84.50% | 5.10% | 1.90% | 8.50% | 53 | 54 | 77701617 |

id = --identity
ovlp = --overlap
letters group identical results
