## Supplementary figure 1 for "Genomic Signatures of Domestication in a Fungus Obligately Farmed by Leafcutter Ants"

**Supplementary Material**

Content:

Supplementary figure S1

Supplementary figure S2

Supplementary table S1

Supplementary table S2

Supplementary table S3

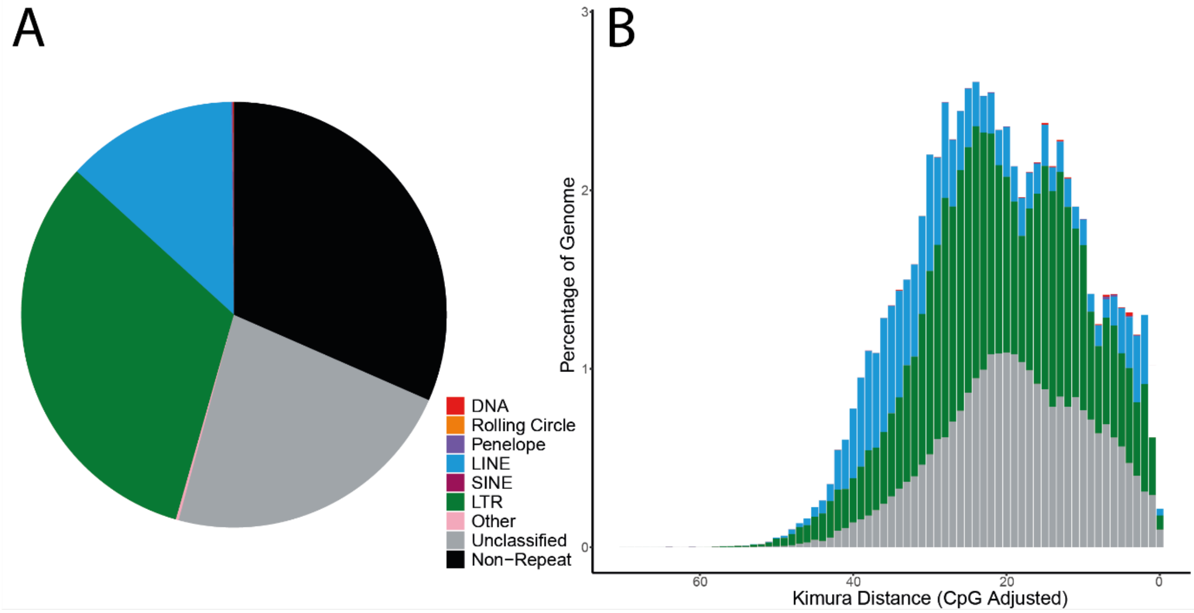


**Figure S1: Transposable element predictions using Earl Grey**. **(A)** The total TE coverage is 67.89% of total genome size and includes 633 distinct TE families. **(B)** The repeat landscape summary shows a general overview of relative TE activity over evolutionary history of the *L. gongylophorus* genome. The reversed X axis displays the Kimura divergence from consensus sequences, indicating TE activity over time—sequences towards the right represent recent activity, while those to the left were active in the past. The Y axis shows the percentage of the genome covered by TEs at varying levels of divergence, revealing periods of increased or decreased TE activity. *Leucoagaricus gongylophorus* shows a notable deficiency in recent TE activity, with two pronounced peaks of past activity at Kimura divergences of approximately 22 and 14. During these peak times, LTR elements dominated expansion, but their activity has since subsided, with low activity observed at a Kimura distance of 0. The observed pattern of intense LTR activity followed by a decline aligns with the burst model of TE behavior, suggesting a cycle of rapid expansion and subsequent suppression by host defense mechanisms, such ach CpG methylation and depletion. The scarcity of recent LTR activity and high CpG deficiency further corroborate this hypothesis.

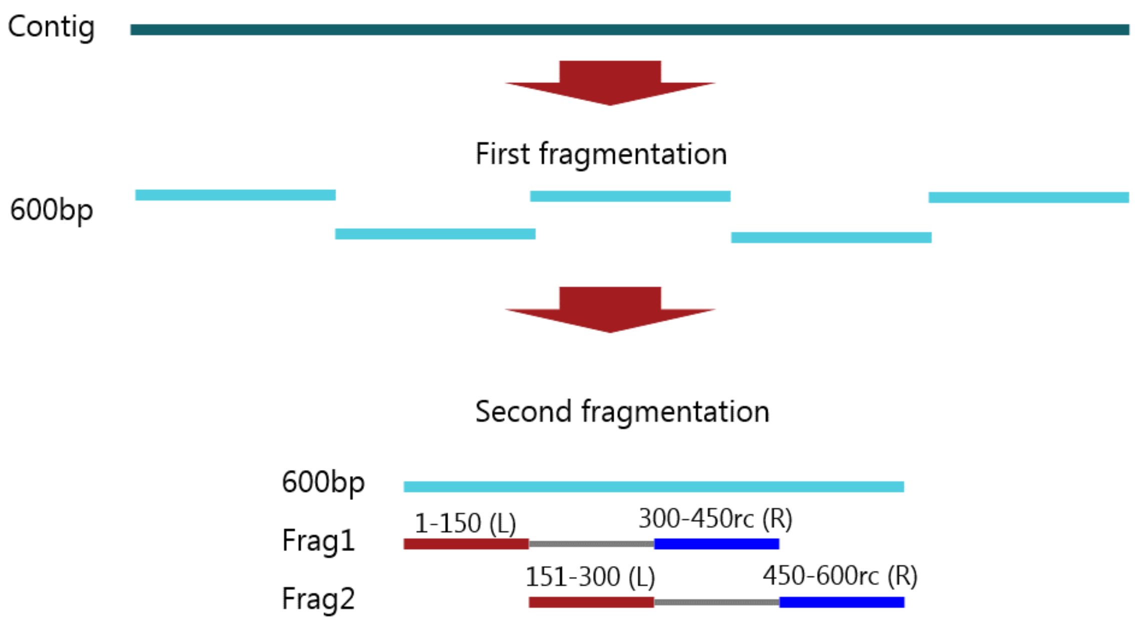


Figure S2: Genome fragmentation steps for Short-Read Simulation (SRS). First, the contigs are first segmented into 600 bp fragments. Next, these 600 bp fragments are divided to form two inserts of 450 bp, with a 300 bp overlap between them. For each insert, the initial 150 bp fragment is designated as the 'left read', and the last 150 bp, which is reverse complement of the original sequence, is designated as the ‘right read’.
